# *Pseudomonas aeruginosa* infection increases palmitoyl carnitine release by host-derived extracellular vesicles

**DOI:** 10.1101/2024.07.13.603378

**Authors:** Rajalakshmy Ayilam Ramachandran, Hamid Baniasadi, Danielle M. Robertson

**Affiliations:** The Departments of Ophthalmology, UT Southwestern Medical Center, Dallas, TX, USA; Biochemistry, UT Southwestern Medical Center, Dallas, TX, USA

## Abstract

*Pseudomonas aeruginosa* (PA), an opportunistic gram-negative pathogen, is the most common pathogen identified in all culture positive cases of infectious keratitis. Extracellular vesicles (EVs) are released by most cells in the body and function in intercellular communication. We have previously reported a change in the proteome of host-derived EVs from corneal epithelial cells during PA infection. In the present study, we investigated changes in the metabolome of host-derived EVs from PA infected (PA-C EVs) and non-infected cells (C EVs). We found that one metabolite, palmitoyl carnitine (PAMC), was significantly upregulated in PA-C EVs. To determine the significance of PAMC release, we investigated the effect of PAMC treatment on corneal epithelial cells and neutrophils. EVs were isolated from culture media using size exclusion chromatography. EVs were then characterized using nanoparticle tracking analysis, transmission electron microscopy, and western blot. Metabolomics was performed using an untargeted approach. We found that palmitoyl carnitine (PAMC) was the most abundant metabolite present in PA-C EVs and was increased more than 3 fold compared to C EVs. Treatment of corneal epithelial cells with increasing levels of PAMC increased nuclear translocation of the NF-κB subunit p65. This was associated with an increase in IL-8 production and neutrophil migration. PAMC also increased levels of mitochondrial calcium. Upon inoculation of corneal epithelial cells with PA, 50 μM PAMC completely eradicated intracellular PA, but stimulated growth of extracellular PA. Taken together, these findings suggest that PA exploits EV release by host cells to deplete PAMC from the intracellular environment.

## Introduction

*Pseudomonas aeruginosa* (PA) is an opportunistic gram-negative pathogen that can cause severe disease in man. In the eye, PA keratitis can lead to vision loss as a result of corneal scarring. PA keratitis has been associated with ocular surface disease, immune compromise, trauma or ocular surgery, and contact lens wear.^1-4^ The pathophysiological mechanisms underlying the disease are complex and multifactorial in nature. However, bacterial virulence factors and an over-exuberant neutrophil response play a role.^5-8^ Invasive strains of PA have been shown to internalize into corneal epithelial cells.^9^ This is mediated in part by lipid-rafts.^10-13^ Indeed, the use lipid depleting agents, such as filipin and methyl-beta-cyclodextrin are associated with reduced levels of intracellular PA.^11, 12^ Intracellular PA has also been shown to induce membrane blebbing in host epithelial cells.^14-17^ This blebbing provides a safe haven for PA to replicate while avoiding clearance by the innate immune system.

Extracellular vesicles (EVs) are membranous nanoscale particles released from all cells in the body and are present in all body fluids.^18^ EVs function as a major mechanism of intercellular communication. The cargo within EVs is derived from their host cell of origin and includes proteins, nucleic acids, and metabolites.^19^ We have recently characterized the proteomic profile of host-derived EVs during corneal infection. We found that EVs released by PA-infected corneal epithelial cells (PA-C EVs) contain both proteins that originate from both the pathogen and host cell.^20, 21^ The inclusion of bacterial proteins in PA-C EVs makes these host-derived vesicles chemotactic for neutrophils. Unlike bacterial-derived EVs however, host-derived EVs do not attenuate the killing capacity of neutrophils.^21^ This sets up an arms race so to speak, whereby bacterial-derived EVs recruit and disable neutrophils, while host-derived EVs (PA-C EVs) function to recruit neutrophils to promote bacterial clearance.

We have recently found that PA impairs host cell mitochondria during invasion.^22^ This was evident by a loss in mitochondrial membrane polarization and the subsequent induction of PTEN-induced kinase 1 (PINK1) and Fun14 domain containing 1 (FUNDC1) mediated mitophagy. This was further associated with an inhibition of the TCA cycle and the electron transport chain, through a targeted reduction in complex I. In that study, we also reported that corneal epithelial cells undergo metabolic rewiring during infection. What remains unknown however, is whether the metabolite profile of EVs is altered during infection. Metabolites are small molecular byproducts and/or intermediates of metabolic pathways. Metabolites can function as signaling molecules and be used by both pathogens and host cells alike. Thus, the EV metabolome may provide insight into the biological pathways that are altered during PA infection.

In the current study, we used an untargeted metabolomics approach to characterize the metabolite profile in PA-C EVs. We found that the metabolite, palmitoyl carnitine (PAMC), was the predominant metabolite that was elevated in EVs during infection. We further characterized the functional role of PAMC in PA invasion and cytokine production. Unexpectedly, we found that PAMC was toxic to intracellular PA following invasion; whereas, PAMC promoted growth of extracellular PA. This finding was restricted to PA in the presence of human corneal epithelial cells. Taken together, these data offer novel insight into the molecular composition of PA-C EVs and have potential translational significance for the eradication of intracellular PA.

## Materials and methods

### Cell culture

A human telomerase-immortalized corneal epithelial (hTCEpi) cell line, developed and characterized by our laboratory, was used in this study.^23^ Cells were cultured in serum-free keratinocyte growth medium with growth factor supplements (KGM, VWR, Randor, PA) and 10% penicillin, streptomycin and amphotericin B (Lonza, Walkersville, MD). The calcium chloride concentration was maintained at 0.15 mM by supplementing additional CaCl_2_ (Calcium Chloride Solution, 0.5 M, VWR, Radnor, PA). For all hTCEpi cell experiments, cells were cultured in media without antibiotics. For the neutrophil experiments, HL-60 cells were purchased from ATCC (Manassas, VA) and routinely cultured in Isocove’s Modified Dulbecco’s Medium (IMDM, ATCC) with 20% fetal bovine serum (FBS, Sigma-Aldrich, St. Louis, MO) and 10% Pen/Strep/Amphotericin B (Lonza). To differentiate cells into neutrophil-like cells, 1.3% dimethyl sulfoxide (DMSO) was added to the cell culture media for 5 days. All cells were maintained at 37°C and 5% CO_2_. For all experiments involving palmitoyl carnitine, palmitoyl-L-carnitine was purchased from Sigma (St. Louis, MO).

### Pseudomonas aeruginosa

A standard invasive strain of *Pseudomonas aeruginosa*, PAO1 (a generous gift from Dr. Suzanne M. J. Fleiszig, UC Berkley) was maintained on tryptic soy agar (TSA, Fisher Scientific, Waltham, MA). Bacteria were cultured overnight (∼14-16 hours) on a TSA slant. Bacteria was resuspended in PBS and adjusted to an OD of ∼1 × 10^8^ CFU/mL. Concentration was measured using a SmartSpec^TM^ Plus spectrophotometer (Bio-Rad, Hercules, CA). The bacterial concentration was confirmed by serial dilution in PBS and plating on tryptic soy agar. All plates were done in triplicate for analysis of colony counts.

### EV isolation

hTCEpi cells were infected with PAO1 at a multiplicity of infection (MOI) of 1 for 6 hours in antibiotic free growth media. 1 × 10^7^ hTCEpi cells were seeded per 150 mm dish. Approximately 60 mLs of culture media were collected and used for each EV isolation. Non-infected hTCEpi cells were used as a control. Cell culture supernatants were centrifuged at 4000Xg for 15 minutes and the resulting pellet was discarded. Culture supernatants were then passed through a 0.2 µm filter and concentrated using 3K concentrators (Pierce protein concentrator PES 3K, Thermo Fisher Scientific, Rockford, IL). EVs were isolated using size exclusion chromatography (qEV columns, Izon Biosciences, Cambridge, MA) according to the manufacturer’s instructions. In brief, 500 μL of concentrated supernatant was passed through the size exclusion column. Fractions were eluted using PBS. After the void volume of 2.7 μL was collected and discarded, the next two 1.5 μL fractions were collected and pooled. Per our recent publication, these two fractions contain EVs with high purity.^21^ EVs from non-infected cells were labeled as C EVs, and EVs from PA infected cells were labeled as PA-C EVs.

### EV characterization

EV protein concentration was measured using a Qubit protein assay (Thermo Fisher Scientific). EV size and concentration were measured using a NanoSight NS300 nanoparticle tracking analyzer (Malvern Panalytical, Worcestershire, UK). The NS300 was configured with a 532 nm laser and a high sensitivity scientific CMOS camera. Each sample was analyzed for 30 seconds, 5 separate times, with a camera level of 15. Data were analyzed using TA 3.4 Build 3.4.4 software. Data from a minimum of 3 independent measurements for each run were used for quantification. For western blotting of known EV markers, EVs were lysed using RIPA buffer and electrophoresed through a 4 - 12% gradient gel (Bio-Rad, Hercules, CA). The proteins were then transferred to polyvinylidene difluoride (PVDF) membranes (Bio-Rad, Hercules, CA). Membranes were then blocked for 1 hour in 5% non-fat milk in PBS containing 0.1% Tween 20 (Bio-Rad, Hercules, CA) at room temperature. Next, membranes were incubated in primary antibodies overnight at 4°C. All primary antibodies were purchased from Abcam (Cambridge, MA). The following antibodies were used: CD9 #ab236630, CD81 #ab109201, CD63 #ab134045, and calnexin #ab112995. Following incubation in a secondary antibody (anti-rabbit HRP #1706515 or anti-mouse HRP #1706516, Bio-Rad, Hercules, CA) for 1 hour, membranes were visualized using enhanced chemiluminescence (Clarity, Bio-Rad, Hercules, CA). Proteins were imaged on an Amersham Imager 600 (Amersham Biosciences, Piscataway, NJ). For morphological analysis of EVs, transmission electron microscopy (TEM) was used. EV samples were mixed 1:1 with 2.5% glutaraldehyde in PBS for 30 minutes. A 400 mesh copper grid with carbon-coated formvar film was incubated with 10 µL of the sample for 20 minutes. After washing with MilliQ water, the grid was stained using 2% uranyl acetate for 10 seconds, followed by another 3 washes with MilliQ water. An FEI Tecnai G2 Spirit Biotwin transmission electron microscope (Thermo Fisher, Waltham, MA) was used to capture EV images. All images were acquired at the Electron Microscopy Core Facility at UT Southwestern Medical Center.

### Metabolomics

To isolate EVs released by PA-infected corneal epithelial cells, hTCEpi cells were first infected with PAO1 at a multiplicity of infection (MOI) of 1 for 6 hours in antibiotic free growth media. Non-infected hTCEpi cells were used as a control. 1 × 10^7^ hTCEpi cells were seeded onto 150 mm dishes. A total of 60 mL of medium per group was collected for EV isolation. Cell culture supernatants were centrifuged at 4000Xg for 15 minutes and the resulting pellet was discarded. EVs were then isolated as described above. Untargeted metabolomics was performed at the Metabolomics Core Facility at UT Southwestern Medical Center.^24^ In brief, 30 μg of EVs were pelleted using the qEV concentration kit (Izon Biosciences, Cambridge, MA) and diluted in 100 μL of ice cold 80% methanol with 5 μM heavy internal standard for 2 hours. All samples were freeze thawed 5 times by alternating for 1 minute each in dry ice and 4^°^C. To collect metabolites, samples were then centrifuged to remove the pellet. The samples with metabolites were filtered using 0.2 µM centrifugal filters and LC-MS was performed using a Sciex QTRAP 6500+ mass spectrometer (Framingham, MA) with EI ion spray ion in both positive and negative mode. A Shimadzu HPLC (Nexera X2 LC-30AD) with analyst 1.7.2 software was coupled to the mass spectrometer (Long Beach, CA). SeQuant® ZIC®-pHILIC 5 μm polymeric 150 × 2.1 mm PEEK coated HPLC column with a temperature of 45^°^C, injection volume of 5 µL and flow rate of 0.15 mL/min was used. A: acetonitrile and solvent B: 20 mM ammonium carbonate with 0.1% ammonium hydroxide and 5 µM of medronic acid. The gradient elution protocol was: 0 min: 80% B, 20 min: 20% B, 20.5 min 80% B, 34 min: 80% for a total of 34 minutes. For the detection and quantification of metabolites, SCIEX MultiQuant 3.0.3 software was used and MetaboAnalyst 5.0 software was used for statistical analysis and data representation. Pathway analysis was performed and a p value of <0.05 and FDR of <0.1 were considered most significant. Metabolites with a fold change of >1.5 and p <0.05 were considered significantly different between C EVs and PA-C EVs.

### Mass spectrometry and proteomic analysis

To isolate EVs released by PA-infected corneal epithelial cells, hTCEpi cells were first infected with PAO1 at a multiplicity of infection (MOI) of 1 for 6 hours in antibiotic free growth media. Non-infected hTCEpi cells were used as a control. 1 × 10^7^ hTCEpi cells were seeded onto 150 mm dishes. A total of 60 mL of medium per group was collected for EV isolation. Cell culture supernatants were centrifuged at 4000Xg for 15 minutes and the resulting pellet was discarded. EVs were isolated as described in the EV isolation section. 7.5 µg of C EVs and PA-C EVs were used for the proteomics studies. All samples were analyzed in triplicate. Mass spectrometry was performed at the Proteomics Core Facility at UT Southwestern Medical Center. Briefly, equal total protein for all experimental groups were subjected to SDS-PAGE. After the samples were run approximately 1 cm into the separating gel, they were cut into pieces and digested overnight with trypsin (Pierce Biotechnology, Rockford, IL) at 37°C. This was followed by reduction with 20 mM dithiothreitol (DTT) at 37°C for 1 hour and alkylation with 27.5 M iodoacetamide in the dark for 20 minutes. After solid-phase extraction cleanup using an Oasis HLB plate (Waters, Milford, MA) the samples were injected into an Orbitrap Fusion Lumos (Thermo, Waltham, MA) mass spectrometer (MS) coupled to an Ultimate 3000 RSLC-Nano liquid chromatography system (Thermo). Samples were eluted as follows: Buffer A was 2% (v/v) ACN and 0.1% formic acid in water, and buffer B was 80% (v/v) ACN, 10% (v/v) trifluoroethanol, and 0.1% formic acid in water. The mass spectrometer was operated in positive ion mode (source voltage of 1.5 kV and an ion transfer tube temperature of 275°C). MS scans were acquired at 120,000 resolutions in the Orbitrap and up to 10 MS/MS spectra at 120,000 resolutions were obtained in the ion trap for each full spectrum using higher-energy collisional dissociation (HCD) for ions with charges 2-7. MS data was analyzed using Proteome Discoverer v2.4 SP1 (Thermo), and Sequest HT was used for peptide identification. Human protein databases were searched in UniProt.

### Lactate dehydrogenase (LDH) assay

To determine whether palmitoyl carnitine was cytotoxic to non-infected corneal epithelial cells, a colorimetric based LDH assay (Abcam, Waltham, MA) was used. hTCEpi cells were seeded onto a 96 well plate at a concentration of 20,000 cells per well in antibiotic free media. Cells were allowed to attach overnight. Cells were then treated with palmitoyl carnitine at a concentration of 5 μM, 50 μM, and 100 μM for 24-72 hours. At the indicated time points, the LDH reaction mix was added to 10 µL of the cell culture supernatant from each well and the absorbance was measured at 450 nm. Data was expressed as percent cytotoxicity. Data is shown from 3-4 independent samples for each run. The experiment was repeated 2 additional times.

### Cytokine ELISAs

hTCEpi cells were seeded in a 96 well plate at a concentration of 20,000 cells per well. To quantify cytokine production by hTCEpi cells, cells were treated with palmitoyl carnitine at a concentration of 5 μM, 50 μM, and 100 μM in media without antibiotics for 24 hours. Supernatants were collected and analyzed using an IL-8 enzyme-linked immunosorbent assay (ELISA, R&D Systems, Minneapolis, MN) according to the manufacturer’s instruction. Briefly, the standards and samples were added into the cytokine antibody pre-coated wells. Unbound samples were washed away and an enzyme-linked polyclonal antibody to detect human IL-8 was added. After washing, the wells were incubated with a substrate against the enzyme and color development was quantified using a BioTek plate reader (Winooski, VT) at 450 nm. Cytokine concentrations were determined using a standard curve. Data is presented from 3 independent biological samples per group. The experiment was repeated an additional 2 times.

### Rhod-2 calcium assay

To quantify levels of mitochondrial calcium, hTCEpi cells were again seeded at a concentration of 20,000 cells per well and allowed to adhere overnight. hTCEpi cells were then treated with palmitoyl carnitine at a concentration of 5 μM, 50 μM, and 100 μM diluted in antibiotic free media. Mitochondrial calcium was measured using a cell permeable calcium indicator, rhod-2 dye. The cells were stained with rhod-2 (Thermo Fisher, Waltham, MA) at a final concentration of 2 µM. A heavy metal-selective chelator, N,N,N′,N′-tetrakis-2-Pyridylmethyl-ethylenediamine (Sigma, St. Louis, MO), was also added at a final concentration of 50 µM to prevent non-specific binding of rhod-2. Following this, cells were treated with one of the three concentrations of PAMC listed above for 30 minutes. Fluorescence was measured using a BioTek plate reader (Winooski, VT) with excitation at 552 nm and emission at 581 nm. Data is presented from 3-4 independent samples. The experiment was repeated 2 additional times.

### NF-κB p65 immunofluorescence

For immunofluorescent studies, 5 × 10^5^ hTCEpi cells were seeded onto 35-mm coverslip bottom dishes (MatTek Corporation, Ashland, MA) and allowed to attach overnight. Cells were then treated with palmitoyl carnitine at a concentration of 5 μM, 50 μM, and 100 μM for 8 hours. Cells were washed with PBS and subsequently fixed in 4% paraformaldehyde (Electron Microscopy Services, Hatfield, PA), and then permeabilized and blocked using PBS containing 0.5% BSA and 0.1% Triton-X 100. Cells were incubated with the primary antibody in PBS containing 0.1% BSA, washed three times with PBS, and then incubated with the secondary antibody containing 0.1% BSA. After washing, cells were mounted using Prolong gold anti-fade reagent containing 4′,6-diamidino-2-phenylindole (DAPI, Invitrogen, Carlsbad, CA). Images were captured using a Leica SP8 laser scanning confocal microscope (Leica Microsystems, Heidelberg, Germany) equipped with a 63× oil objective. The primary antibody that was used was NF-κB p65 #8242 (Cell Signaling Technology, Danvers, MA). The secondary antibody that was used was an anti-rabbit Alexa 488 secondary antibody #A21206 (Thermo Fisher Scientific, Rockford, IL). Staining was visualized using a 488 nm excitation laser (NF-κB p65) and a UV laser (DAPI). Images were sequentially scanned to avoid spectral crosstalk.

### Bacterial quantification

To quantify bacteria, 1 × 10^5^ cells were seeded onto 12 well plates with a total volume of 1 mL media without antibiotics overnight. hTCEpi cells were treated with 5 μM or 50 μM of palmitoyl carnitine for 24 hours to assess the total bacterial growth (intra and extracellular). At the end of that time period, samples were inoculated with 40 µL of PA at a concentration of 1 × 10^6^ CFU/mL and incubated for an additional 2 hours. For total bacteria (comprised of both extra- and intracellular bacteria), the cell culture supernatant was collected. The cells were then lysed in 0.25% triton X 100 and lysates were combined with the cell culture supernatant. Samples were then serially diluted in PBS and plated on tryptic soy agar plates for enumeration (Fisher Scientific, Waltham, MA). To assess the effect palmitoyl carnitine on the growth of PA in the absence of corneal epithelial cells, 12 well plates were filled with 1 mL media without antibiotics containing 5 μM or 50 μM of palmitoyl carnitine for 24 hours. At the end of the incubation period, samples were inoculated with 40 µL of PA with a concentration of 1 × 10^6^ CFU/mL and incubated for an additional 2 hours. To quantify the bacterial load, the sample was serially diluted in PBS and plated on tryptic soy agar plates for enumeration (Fisher Scientific, Waltham, MA). All samples were plated in triplicate. The experiment was repeated an addition 2 times.

### Gentamicin protection assay

To quantify only intracellular bacteria, 5 x 10^5^ hTCEpi cells were seeded onto a 6 well plate and allowed to adhere overnight. Cells were then treated with 5 μM or 50 μM palmitoyl carnitine for 24 hours. At the 24 hour time point, cells were inoculated with PA for 2 hours. The bacterial concentration was approximately ∼500 bacteria per 1 hTCEpi cell. At the 2 hour time point, gentamicin (200 µg/mL, Sigma, St. Louis, MO) was added to kill all extracellular bacteria for either 1 hour or 6 hours. The treated cells were lysed using 0.25% Triton X 100 for 15 minutes at room temperature to release intracellular bacteria. Lysates were serially diluted in PBS and plated onto tryptic soy agar. All samples were plated in triplicate. The entire experiment was repeated 2 additional times.

### Neutrophil chemotaxis

Neutrophil chemotaxis was assessed using a Boyden chamber fluorometric assay (Cell Biolabs, San Diego, CA). To generate the cell culture supernatants used in this assay, 20,000 hTCEpi cells were seeded onto 96 well plates and allowed to adhere overnight. The cultured cells were treated for 24 hours in media containing 5μM, 50 μM, and 100 μM palmitoyl carnitine. Supernatants were collected for use in the Boyden chamber. Briefly, 100 μL of IMDM containing 5 × 10^5^ DMSO differentiated HL-60 cells were seeded onto the top chamber and 150 μL of hTCEpi cell culture supernatants in the bottom chamber. After 2 hours, the migrated cells in the bottom chamber were lysed and fluorescence was measured using a BioTek plate reader (Winooski, VT). All samples were performed in triplicate and each experiment was repeated a minimum of two additional times.

### Neutrophil respiratory burst

Neutrophil respiratory burst was measured using a dihydrorhodamine 123 fluorescent assay in a 96-well plate (Abcam, Waltham, MA). Briefly, 1 × 10^5^ DMSO-differentiated HL-60 cells were stained with dihydrorhodamine 123 and incubated along with 5 μM, 50 μM, and 100 μM palmitoyl carnitine in IMDM for 1, 2 or 3 hours. The fluorescence was measured using a BioTek plate reader (Winooski, VT) with excitation at 552 nm and emission at 581 nm. All samples were plated in triplicate and each experiment was repeated an additional two times.

### Neutrophil bacteria killing

Neutrophil bacteria killing was assessed by treating 1 × 10^5^ DMSO-differentiated HL-60 cells with 5 μM, 50 μM, and 100 μM palmitoyl carnitine in a 6 well plate. PAO1 was added at a concentration of 5 × 10^4^ bacteria per well. We used IMDM with 10% human serum as the reaction media. The cells were incubated for 2 hours at 37^°^C. Following that, the total viable bacterial count in each experimental group was determined by serial dilution in PBS and plating on tryptic soy agar. All samples were plated in triplicate and each experiment was repeated at least 2 additional times.

### Statistics

All data are presented as mean ± standard deviation. A t-test was used to determine differences between 2 groups. For multiple groups, we used a one-way ANOVA with a Tukey’s post hoc multiple comparison test. A P <0.05 was considered statistically significant.

## Results

### EV isolation and characterization

EVs were isolated from cell culture supernatants using qEV columns and characterized by size using nanoparticle tracking analysis. In agreement with our prior studies, EVs from F1 and F2 for both PA infected and non-infected hTCEpi cells had particles in the size range of ∼100-200 nm (Figure 1A). These fractions (F1 and F2) were combined as the EV fraction and examined using TEM. TEM confirmed the presence of EVs in both groups (Figure 1B). Western blot was then performed for known EV markers. The tetraspanin proteins CD9, CD81, and CD63 were all enriched in EVs compared to whole cell lysates. The protein calnexin, an endoplasmic reticulum protein that is not present in EVs, was absent in the EVs but detected in whole cell lysate (Figure 1C). Together, these findings confirmed the successful isolation of EVs.

**Figure 1:**
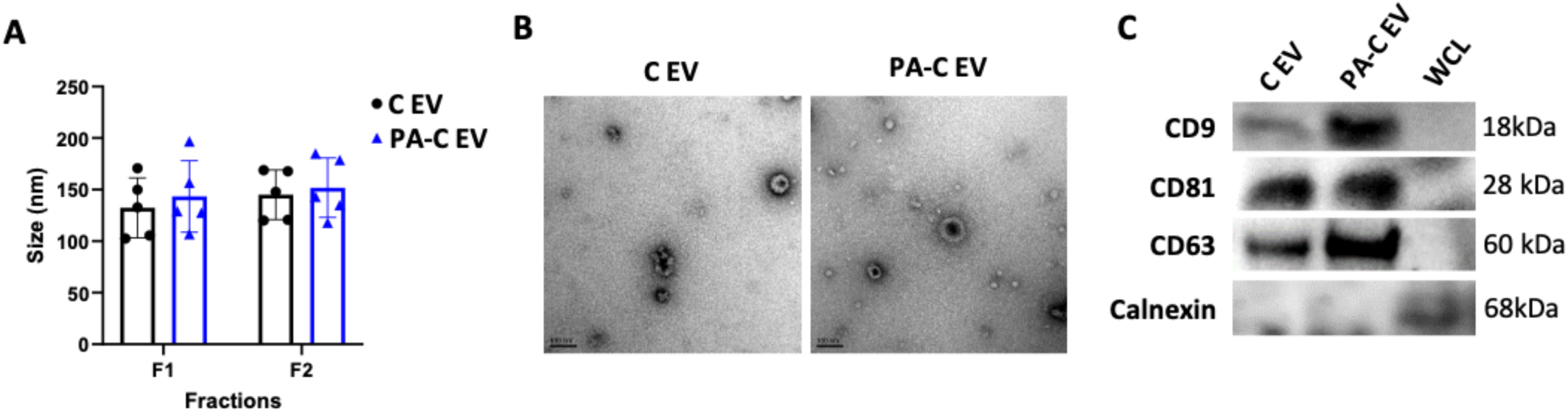
EV characterization. EVs were isolated from non-infected and PAO1 infected hTCEpi cells by size exclusion chromatography. (A) NTA analysis of EV fractions, F1 and F2, showed no difference in EV size between non-infected and infected groups. (B) TEM images confirmed the presence of EVs smaller than 200 nm. Scale bar 100 nm. (C) Western blot for EV markers CD9, CD81, and CD63. Calnexin, which is not present in EVs, was only present in the whole cell lysate (WCL). Data presented as mean ± standard deviation, N≥3, t–test.

### PA infected corneal epithelial cell-derived EVs are enriched in palmitoyl carnitine

Using an untargeted metabolomics approach, we assayed the metabolites present in C EVs and PA-C EVs. We identified a total of 30 metabolites in both groups of EVs (Figure 2A). Subsequent pathway analysis of these metabolites was then performed. The most significant pathways included alanine, aspartate and glutamate metabolism; butanoate and propanoate metabolism; and phenylalanine, tyrosine and tryptophan biosynthesis (Figure 2B). Of the 30 metabolites, palmitoyl carnitine and glycerate were significantly upregulated in PA-C EVs, while cortisol and aniline-2-sulfonate were significantly downregulated compared to C EVs (Figure 3, A-D). Of the two metabolites that were increased, palmitoyl carnitine was increased over 3-fold in PA-C EVs. Pathway analysis of the significantly altered metabolites in the PA-C EV group were related to glycerolipid metabolism and the pentose phosphate pathway (Figure 3E).

**Figure 2:**
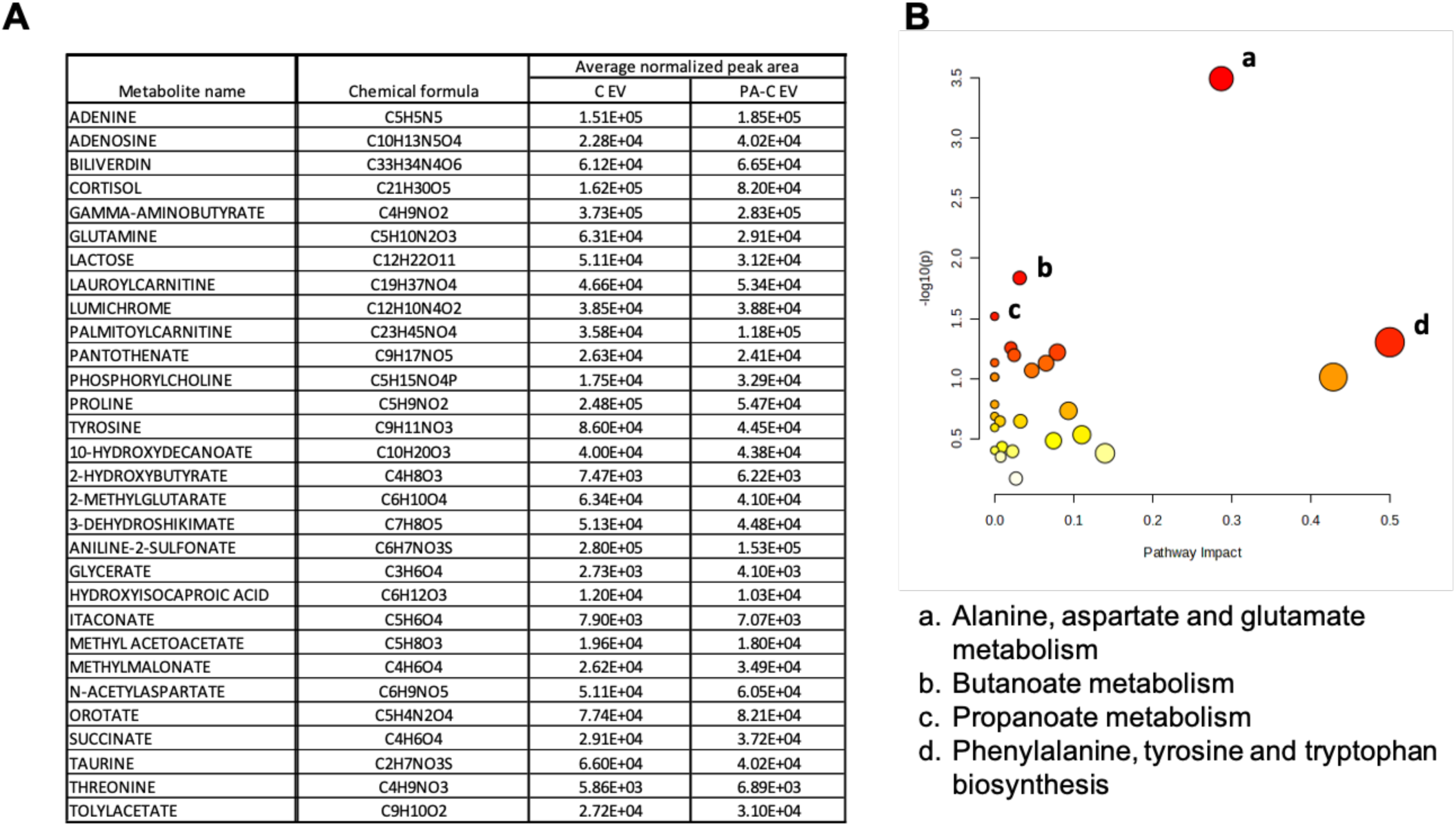
Metabolites profile of C EVs and PA-C EVs. (A) List of all metabolites present in C EVs and PA-C EVs. (B) Pathway impact analysis of the most significant metabolite pathways in C EVs and PA-C EVs. Untargeted metabolomics N= 3, MetaboAnalyst 6.0.

**Figure 3:**
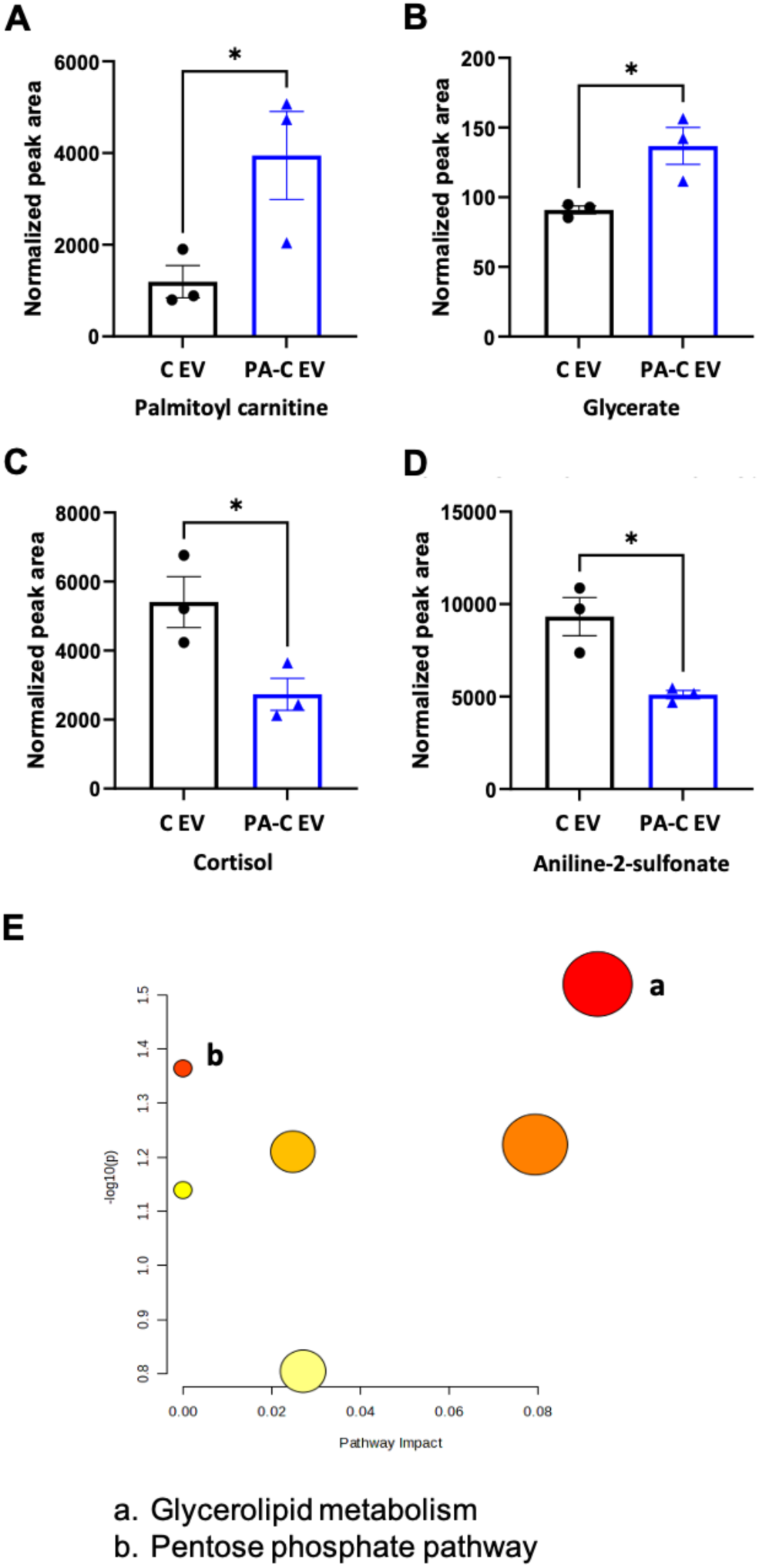
Four metabolites differed between C EVs and PA-C EVs. The peak areas detected by LC-MS were normalized using total protein for comparison between the two groups. (A-B) Palmitoyl carnitine (A) and glycerate (B) were significantly increased in PA-C EVs. (C-D) Cortisol (C) and aniline-2-sulfonate (D) were significantly decreased in PA-C EVs. (E) Pathway impact analysis of the pathways that were significantly altered in PA-C EVs compared to C EVs. Data presented as mean ± standard deviation, N=3, t-test, *p<0.05. Untargeted metabolomics, MetaboAnalyst 6.0.

### Carnitine pathway associated metabolites and enzymes were decreased in corneal epithelial cells during PA infection

Since palmitoyl carnitine was significantly enriched in PA-C EVs, we next explored the related pathway metabolites and enzymes. Using proteomics, we found that the enzymes carnitine O-palmitoyl transferase 2 and carnitine O-acyl transferase were reduced in PA-C EVs (Supporting figure 1, A&B). We also performed untargeted metabolomics on corneal epithelial cells during PA infection. We found that the metabolites related to the carnitine pathway, such as O-acyl carnitine and deoxy carnitine, were significantly decreased. L-carnitine however, was unchanged (Figure 4, A-C). These data suggest that carnitine metabolism is downregulated in corneal epithelial cells during PA infection. To confirm that these changes were not due to cytotoxicity, we next performed a lactate dehydrogenase assay. Cells were exposed to varying concentrations of PAMC ranging from 5-100 µM after 24, 48, and 72 hours. Cytotoxicity was only noted at the highest concentration tested, 100 µM PAMC. This resulted in hTCEpi cell cytotoxicity of 54% at 24 hours, 6% at 48 hours, and 31% at 72 hours (Supporting figure 2, A-C). There was no cytotoxic effect of 5 or 50 µM PAMC on hTCEpi cells at any of the time points tested.

**Figure 4:**
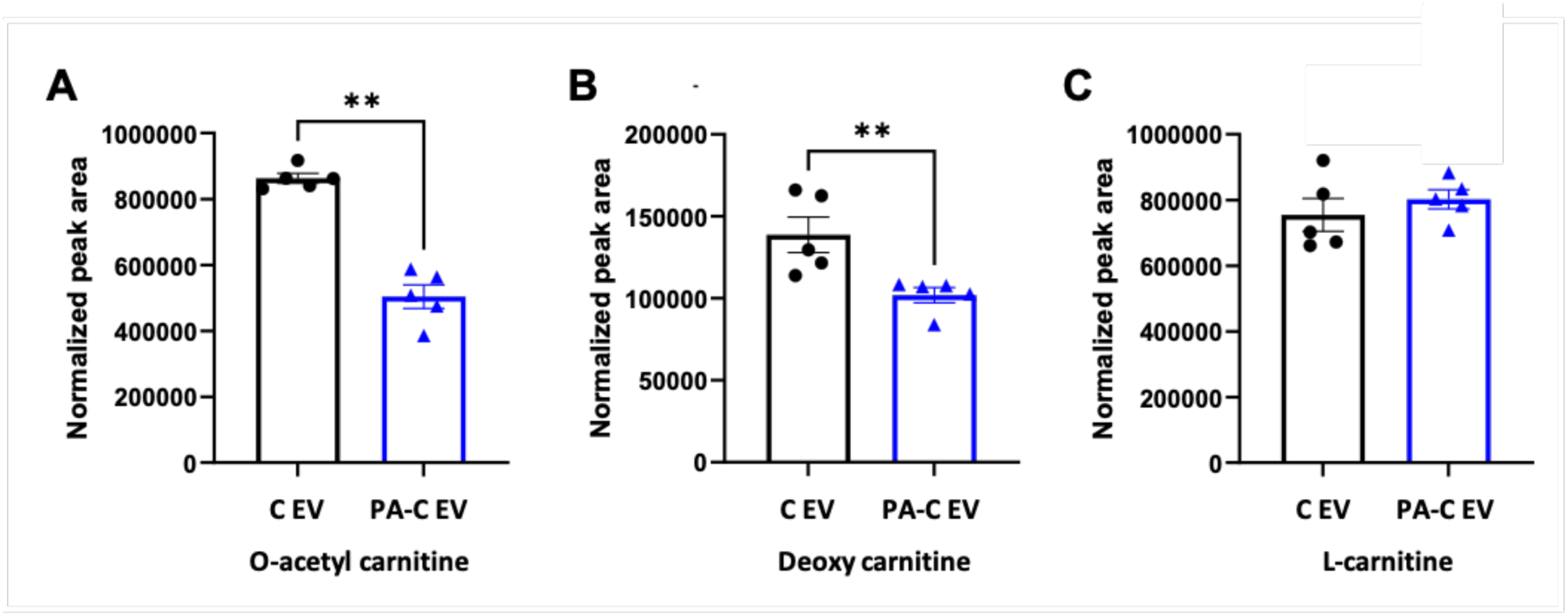
Carnitine metabolites were decreased in PAO1 infected hTCEpi cells. The peak areas detected by LC-MS were normalized using total protein for comparison between the two groups. (A-B) O-acetyl carnitine (A) and deoxy carnitine (B) were decreased in infected cells. (C) L-carnitine was unchanged during infection. Data represented as mean ± standard deviation, N=3, t-test, **p<0.01.

### Palmitoyl carnitine induced a pro-inflammatory cytokine response through activation of NF-κB

Prior studies have shown that PAMC is capable of inducing a pro-inflammatory cytokine response in cancer cells.^25^ To test the effects of PAMC on cytokine production in hTCEpi cells, we treated cells with 5 μM, 50 μM, or 100 μM PAMC for 8 hours. Immunofluorescent staining demonstrated the nuclear translocation of the p65 subunit of NF-κB, a transcription factor that regulates cytokine production (Figure 5A). Nuclear levels of p65 increased in a dose dependent manner, with 100 μM showing complete translocation from the cytoplasm to the nucleus. We next tested the effects of PAMC on the pro-inflammatory cytokine response by quantifying IL-6 and IL-8 release in cell culture supernatants. PAMC was not capable of inducing IL-6 secretion in hTCEpi cells (data not shown). However, 24 hours of exposure to PAMC induced IL-8 secretion in hTCEpi cells in a concentration independent manner (Figure 5B).

**Figure 5:**
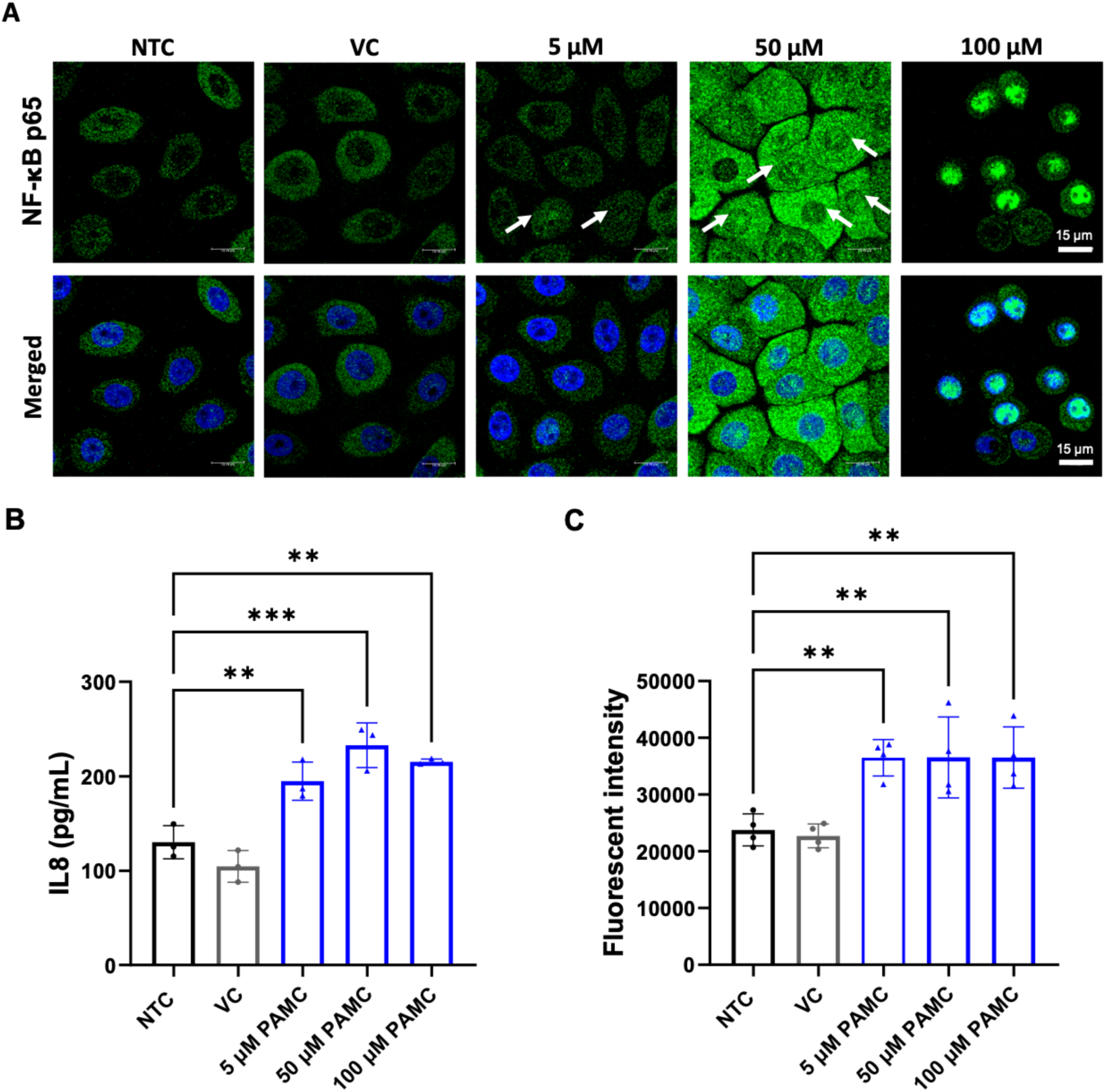
PAMC triggered inflammation in non-infected hTCEpi cells. (A) Immuno-fluorescence for NF-κB p65. Nuclei were labeled with DAPI. Eight hours of treatment with PAMC increased nuclear translocation of p65 in a concentration dependent manner. Arrows indicate nuclear p65. Images representative of 3 repeated experiments, scale bar: 15 μM. (B) IL-8 levels in hTCEpi cell culture supernatants after 24 hours of treatment with PAMC. (C) Neutrophil migration was detected using a Boyden-Chamber assay. Cell culture supernatants from PAMC treated cells induced neutrophil migration. Data represented as mean ± standard deviation, N≥3, one-way ANOVA with Tukey’s post hoc multiple comparison test, **p<0.01, ***p<0.001. VC: vehicle control (DMSO), NTC: no template control.

Since IL-8 is a major chemotactic factor for neutrophils, we next tested the ability of cell culture supernatants from PAMC-treated cells to stimulate neutrophil migration.^26, 27^ Neutrophil migration was quantified using a Boyden chamber assay. hTCEpi cells were treated with 5 µM, 50 μM, or 100 μM PAMC for 24 hours. Supernatants from all PAMC treated cultures were able to induce significant neutrophil migration in a concentration independent manner compared to the non-treated control (Figure 5C). This data suggests that PAMC is capable of promoting the innate immune response in corneal epithelial cells. To further investigate the effect of PAMC on neutrophils, we next quantified neutrophil respiratory burst. DMSO-differentiated HL-60 cells were exposed to PAMC for 1, 2 or 3 hours. As shown in Figure 6 (A-C), there was a dose dependent increase in respiratory burst at the 2 hour and 3 hour time points. Despite the increase in respiratory burst, PAMC treatment had no effect on bacterial killing (Figure 6D).

**Figure 6:**
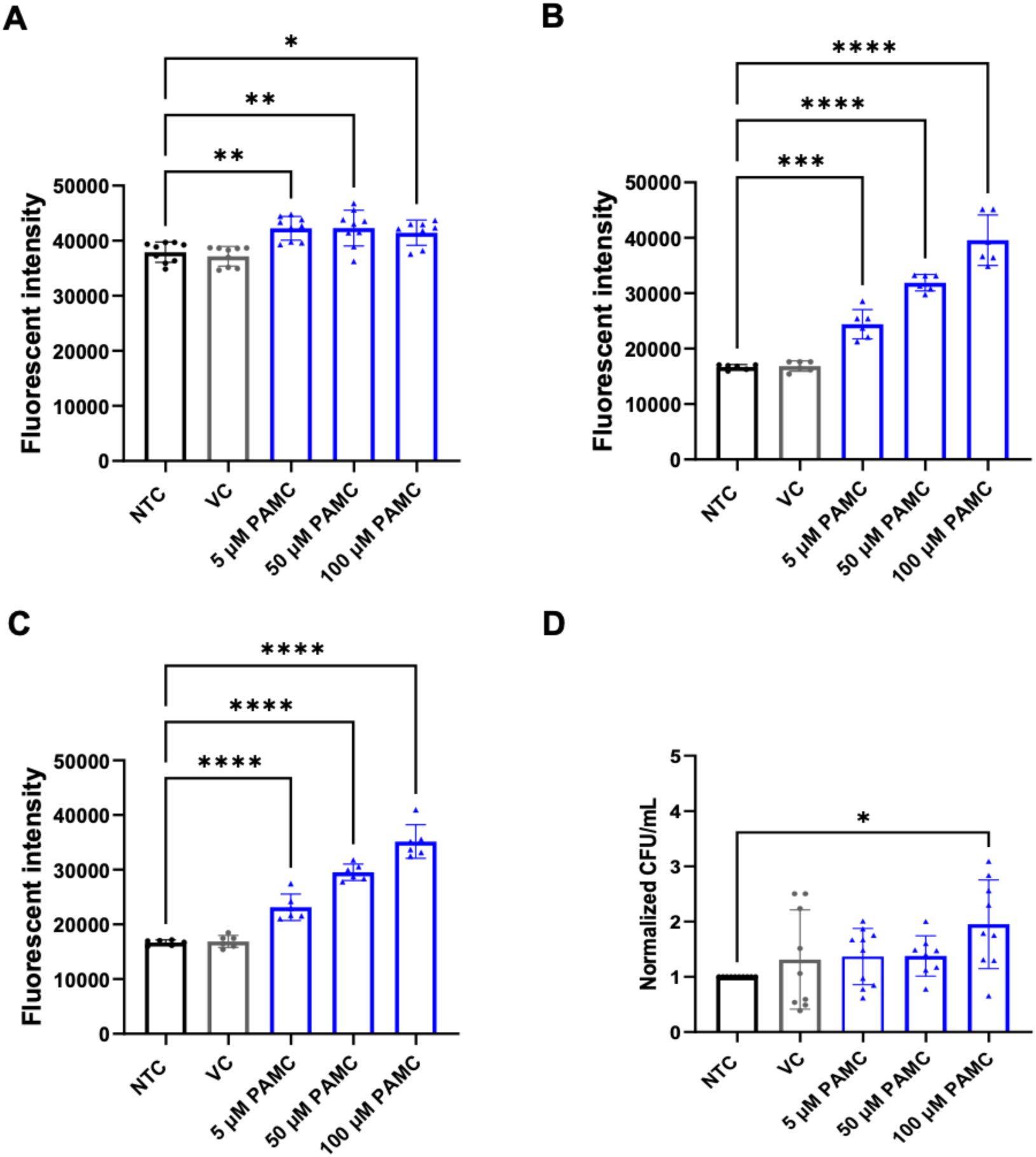
Palmitoyl carnitine stimulated neutrophil respiratory burst. DMSO differentiated neutrophils were treated with PAMC to quantify neutrophil burst. (A-C) PAMC treatment for 1 hour (A), 2 hours (B), and (C) 3 hours. (D) Despite the increase in respiratory burst, there was no increase in bacterial killing by PAMC-treated neutrophils. CFU data normalized to the control. Data represented as mean ± standard deviation, N≥3, one-way ANOVA with Tukey’s post hoc multiple comparison test, *p<0.05, **p<0.01, ***p<0.001, and ****p<0.0001. VC: vehicle control (DMSO), NTC: no template control.

### Mitochondrial calcium increased in response to palmitoyl carnitine

Control of mitochondrial calcium is crucial for maintaining ATP production and cellular respiration.^28^ In a prior study, PAMC was shown to induce calcium influx in prostate cancer cells.^25^ Here, we looked at mitochondrial calcium influx in corneal epithelial cells upon PAMC exposure. Similarly, we found that PAMC increased mitochondrial calcium levels after 30 minutes (Figure 7). Since 50 μM PAMC is not toxic to corneal epithelial cells, this influx in mitochondrial calcium may be sufficient to drive increased mitochondrial respiration. Whereas, 100 μM, which is associated with cell toxicity, may push mitochondrial calcium levels to a toxic level. These findings require clarification in future studies.

**Figure 7:**
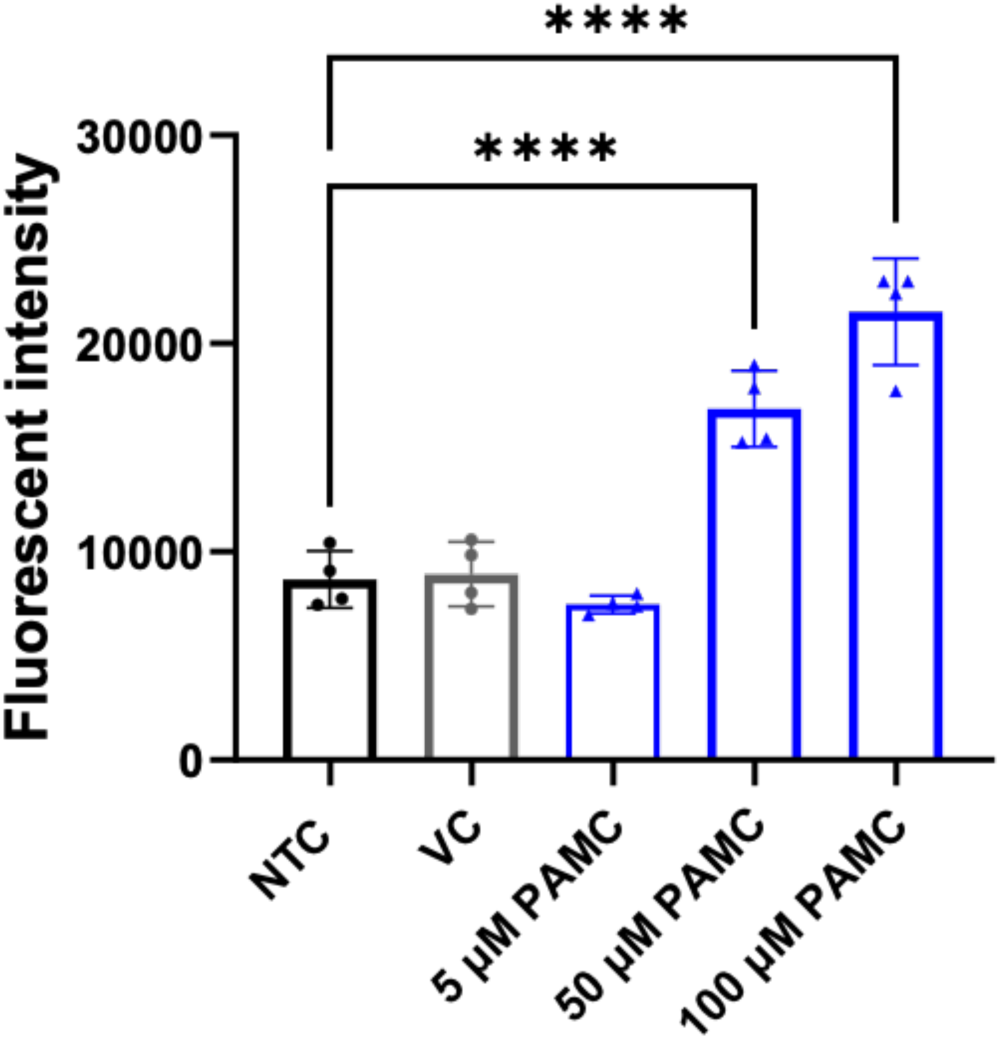
Palmitoyl carnitine increased mitochondrial calcium in hTCEpi cells. 50 μM and 100 μM PAMC increased mitochondrial calcium levels. Data presented as mean ± standard deviation, N≥3, one-way ANOVA with Tukey’s post hoc multiple comparison test, *p<0.05, **p<0.01, ***p<0.001, and ****p<0.0001. VC: vehicle control (DMSO), NTC: no template control.

### Palmitoyl carnitine differentially promotes and inhibits PA growth

To investigate the effects of PAMC on PA, we initially tested the effect of PAMC on planktonic culture of PA in the presence of our standard growth medium. To do this, we directly inoculated PA into growth medium containing PAMC. At 5 µM and 50 µM concentrations, PAMC had no effect on the growth of PA (Figure 8, A-B). However, when PAMC was added to hTCEpi cell monolayers cultured in the same growth media at a concentration of 5 μM, the amount of PA increased significantly. We saw a similar effect for 50 µM PAMC (Figure 8, C-D). This indicates that changes in PA following contact with host cells altered its response to PAMC. This finding may have potential implications in the contact lens wearing eye, where reduced tear clearance leads to the accumulation of cellular byproducts in the microenvironment under the lens.

**Figure 8:**
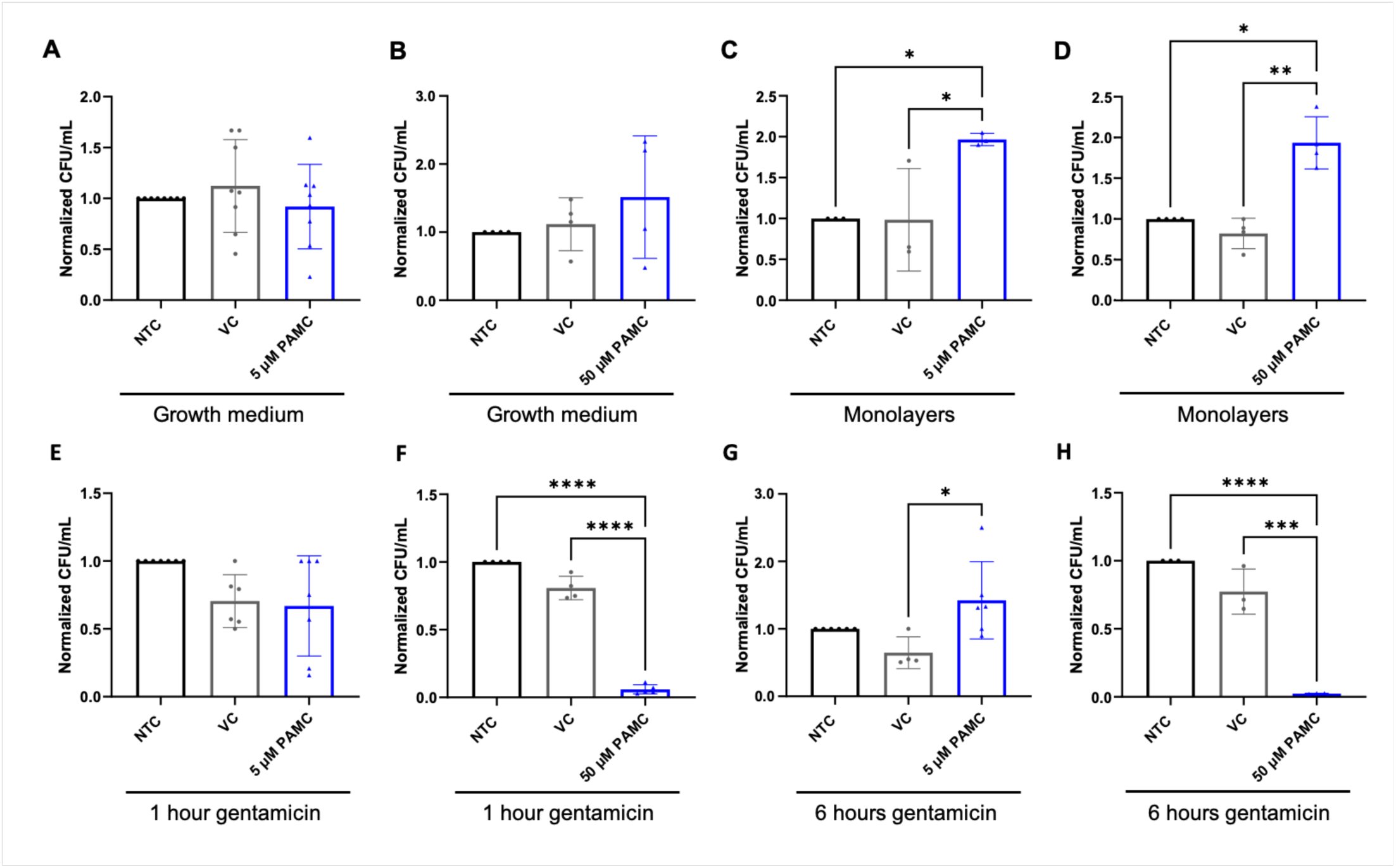
PAMC inhibited intracellular bacterial replication in hTCEpi cells. (A-D) PAO1 was cultured for 2 hours either in culture medium without hTCEpi cells or added to hTCEpi cell monolayers. Both conditions were pretreated with PAMC for 24 hours prior to inoculation. (A-B) PAMC had no effect on levels of PAO1 in growth medium. (C-D) In PA-infected cell culture monolayers, PAMC treatment increased in levels of PAO1 compared to controls. (E-F) Two hours after PA infection, cells were treated with 200 μg/mL gentamicin to kill extracellular PA for 1 hour. While treatment with 5 μM PAMC had no effect on levels of intracellular PA (E), treatment with 50 μM PAMC reduced intracellular levels of PA to almost undetectable levels (F). (G-H) Two hours after PA infection, cells were treated with 200 μg/mL gentamicin to kill extracellular PA for 1 hour. Cultures were then washed and replenished with fresh medium containing gentamicin for an additional 6 hours. Similar to the 1 hour treatment time, 5 μM PAMC had no effect on levels of intracellular PA (G). No bacteria were detected after treatment with 50 μM PAMC for 6 hours. Data are represented as mean ± standard deviation, N≥3. In all experiments, plate counts were normalized to the no treatment control. VC: vehicle control (DMSO), NTC: no template control. One-way ANOVA with Tukey’s post hoc multiple comparison test, *p<0.05, **p<0.01, ***p<0.001, and ****p<0.0002.

We next examined the effect of PAMC on intracellular growth. Again, cell culture monolayers were treated with PAMC for 24 hours prior to infection with PA. Two hours post infection, inoculated cells were then treated with gentamicin for 1 hour to kill all of the extracellular PA. At the 5 μM concentration of PAMC, there was no effect on intracellular levels of PA (Figure 8E). However, at 50 μM of PAMC, there was a significant reduction in PA (Figure 8F). This suggests that PAMC may inhibit the invasion of PA into corneal epithelial cells. We next sought to determine whether PAMC inhibited PA replication. Here, we added gentamicin for 1 hour to kill all of the extracellular PA. After 1 hour, cells were washed and media containing gentamicin was added for an additional 5 hours. Similar to the invasion assay, 5 μM PAMC had no effect on intracellular levels of PA (Figure 8G). In cells treated with 50 μM PAMC however, no intracellular PA was detected (Figure 8H). Taken together, these data suggest that PAMC is able to either prevent invasion and/or kill intracellular PA. Lastly, we wanted to determine whether a shorter treatment with PAMC had the same effect as 24 hours. Here, cells were treated with 50 μM PAMC for 2 hours. At this higher concentration, 2 hours of PAMC again reduced intracellular levels of PA to almost undetectable levels (Supplementary figure 3).

## Discussion

Here we show that PA infection alters the metabolite profile of EVs released by corneal epithelial cells. The most notable effect was the upregulation of PAMC. This effect appears to be specific to PAMC, since O-acetyl-carnitine and deoxy carnitine were decreased and L-carnitine was unchanged. PAMC is an acyl-carnitine that functions as a key intermediate in the carnitine shuttle. There it serves as the temporary form of palmitic acid that is able to traverse the inner mitochondrial membrane into the mitochondrial matrix via carnitine-acylcarnitine translocase. Once in the mitochondrial matrix, PAMC is then converted to palmitoyl-CoA by carnitine O-palmitoyl transferase 2 (CPT2). Consistent with the export of PAMC via EV release, proteomics also showed a decrease in CPT2 in response to PA infection. The reduction in this enzyme alone would prevent the conversion of PAMC to palmitoyl-CoA. Together, these data suggest that PA infection of corneal epithelial cells impairs the beta-oxidation of fatty acids at this early time point.

PA has been shown to hydrolyze acyl-carnitines such as PAMC into carnitine and a free fatty acid.^29^ Carnitine has also been shown to protect PA against osmotic stress, induce secretion of the virulence factor phospholipase C, and provide carbon and nitrogen as a source of nutrition.^30, 31^ However, PA can neither synthesize carnitines nor acyl-carnitines and it must acquire them from the environment.^29^ Thus, the upregulation of PAMC in corneal epithelial cell-derived EVs may function as a protective mechanism to prevent invasive PA from scavenging carnitine within infected epithelial cells. Further research is needed to investigate this finding.

PAMC may also play a role in inflammatory signaling. Indeed, our data shows a dose-dependent relationship between PAMC and nuclear translocation of the NF-κB subunit p65. The activation of the NF-κB pathway is known to trigger the upregulation of many pro-inflammatory cytokines and chemokines, leading to immune cell recruitment and activation.^32^ In our model, we found a significant increase in IL-8 in cell culture supernatants following PAMC treatment. In concert with this, PAMC induced neutrophil migration and respiratory burst. Given that IL-8 is a major neutrophil chemokine, it is not surprising that this increase promoted neutrophil migration *in vitro*. IL-8 has also been shown to be necessary for neutrophil priming for oxidative burst, which is similarly increased by PAMC.^33^ Together, it appears that PAMC serves as a pro-inflammatory metabolite that induces NF-κB activation, which in turn leads to the subsequent secretion of IL-8 and neutrophil recruitment. While the above processes are necessary for the innate immune response to PA, excessive inflammation and persistent recruitment of immune cells in the cornea is a major cause of severe tissue damage.

Mitochondrial dysfunction has been associated with disease in many cells and tissues. We have previously shown that both hyperglycemic and hypersomolar stress result in mitochondria dysfunction in the corneal epithelium.^34-36^ More recently, we have reported on the ability of PA to induce mitochondrial dysfunction by targeting complex 1 of the electron transfer chain. This is accompanied by the induction of mitophagy and metabolic rewiring to support the host cell. The observed impairment in the beta oxidation of fatty acids is in line with our prior work. Moreover, it appears that the downregulation of this pathway is sufficient to trigger the release of PAMC via EVs into the extracellular space. Whether enough PAMC is then taken up by host cells to promote an overall increase in mitochondrial calcium levels remains to be determined. Calcium regulation within mitochondria is extremely complex. While a moderate increase in mitochondrial calcium is able to enhance mitochondrial function, excessive levels of mitochondrial calcium drive the induction of cell death.

The most unexpected finding in this study was the ability of PAMC to reduce levels of intracellular PA to non-detectable levels. This finding was restricted to intracellular bacteria. In contrast, the total bacterial count was increased for extracellular bacteria, but only in the presence of corneal epithelial cells. This suggests that cell-mediated changes in PA enhances the ability of the bacteria to slurp up PAMC and use it to promote bacterial metabolism and replication. PA has a highly branched respiratory chain and is able to use oxygen and nitrogen as terminal electron acceptors. This allows PA to quickly switch from aerobic to anaerobic metabolism. Our prior work has shown that oxygen consumption by PA appears to change depending on whether the bacteria is intracellular or extracellular.^22^ This raises the question of whether PAMC exerts a toxic effect on anaerobic PA due to this metabolic shift. If this proves to be true, it means that PA is able to exploit the host EV release pathway to expel a potentially cytotoxic metabolic. Additional studies are needed to investigate this new hypothesis.

Overall, this study demonstrates that PAMC released through EVs during PA infection may function as a double-edged sword in regulating disease pathogenesis. This has important implications for contact lens wear. For extracellular bacteria, the presence of EVs containing PAMC that accumulate under the lens may lead to increased bacterial growth. In contrast to this, the ability to eradicate PA from the intracellular environment using a known metabolite without any corresponding cytotoxicity may have therapeutic potential. Metabolites have been shown to have antimicrobial potential and, in some cases, to mediate antibiotic resistance. Thus, significant work is needed to delve into the role of PAMC in mediating bacterial survival and replication in the corneal epithelium, with a specific focus on metabolic changes in the pathogen.

## References

[1] Poggio EC, Glynn RJ, Schein OD, Seddon JM, Shannon MJ, Scardino VA, Kenyon KR: The incidence of ulcerative keratitis among users of daily-wear and extended-wear soft contact lenses. N Engl J Med 1989, 321:779–83.

[2] Schein OD, Glynn RJ, Poggio EC, Seddon JM, Kenyon KR: The relative risk of ulcerative keratitis among users of daily-wear and extended-wear soft contact lenses. A case-control study. Microbial Keratitis Study Group. N Engl J Med 1989, 321:773–8.

[3] Stapleton F, Naduvilath T, Keay L, Radford C, Dart J, Edwards K, Carnt N, Minassian D, Holden B: Risk factors and causative organisms in microbial keratitis in daily disposable contact lens wear. PLoS One 2017, 12:e0181343.

[4] Pachigolla G, Blomquist P, Cavanagh HD: Microbial keratitis pathogens and antibiotic susceptibilities: a 5-year review of cases at an urban county hospital in north Texas. Eye Contact Lens 2007, 33:45–9.

[5] Steuhl KP, Döring G, Henni A, Thiel HJ, Botzenhart K: Relevance of host-derived and bacterial factors in Pseudomonas aeruginosa corneal infections. Invest Ophthalmol Vis Sci 1987, 28:1559–68.

[6] Robertson DM, Parks QM, Young RL, Kret J, Poch KR, Malcolm KC, Nichols DP, Nichols M, Zhu M, Cavanagh HD, Nick JA: Disruption of contact lens-associated Pseudomonas aeruginosa biofilms formed in the presence of neutrophils. Invest Ophthalmol Vis Sci 2011, 52:2844–50.

[7] Robertson DM, Rogers NA, Petroll WM, Zhu M: Second harmonic generation imaging of corneal stroma after infection by Pseudomonas aeruginosa. Sci Rep 2017, 7:46116.

[8] Hinojosa JA, Patel NB, Zhu M, Robertson DM: Antimicrobial Efficacy of Contact Lens Care Solutions Against Neutrophil-Enhanced Bacterial Biofilms. Transl Vis Sci Technol 2017, 6:11.

[9] Fleiszig SM, Zaidi TS, Fletcher EL, Preston MJ, Pier GB: Pseudomonas aeruginosa invades corneal epithelial cells during experimental infection. Infect Immun 1994, 62:3485–93.

[10] Yamamoto N, Yamamoto N, Jester JV, Petroll WM, Cavanagh HD: Prolonged hypoxia induces lipid raft formation and increases Pseudomonas internalization in vivo after contact lens wear and lid closure. Eye Contact Lens 2006, 32:114–20.

[11] Yamamoto N, Yamamoto N, Petroll MW, Cavanagh HD, Jester JV: Internalization of Pseudomonas aeruginosa is mediated by lipid rafts in contact lens-wearing rabbit and cultured human corneal epithelial cells. Invest Ophthalmol Vis Sci 2005, 46:1348–55.

[12] Yamamoto N, Yamamoto N, Petroll MW, Jester JV, Cavanagh HD: Regulation of Pseudomonas aeruginosa internalization after contact lens wear in vivo and in serum-free culture by ocular surface cells. Invest Ophthalmol Vis Sci 2006, 47:3430–40.

[13] Zaidi T, Bajmoczi M, Zaidi T, Golan DE, Pier GB: Disruption of CFTR-dependent lipid rafts reduces bacterial levels and corneal disease in a murine model of Pseudomonas aeruginosa keratitis. Invest Ophthalmol Vis Sci 2008, 49:1000–9.

[14] Angus AA, Evans DJ, Barbieri JT, Fleiszig SM: The ADP-ribosylation domain of Pseudomonas aeruginosa ExoS is required for membrane bleb niche formation and bacterial survival within epithelial cells. Infect Immun 2010, 78:4500–10.

[15] Angus AA, Lee AA, Augustin DK, Lee EJ, Evans DJ, Fleiszig SM: Pseudomonas aeruginosa induces membrane blebs in epithelial cells, which are utilized as a niche for intracellular replication and motility. Infect Immun 2008, 76:1992–2001.

[16] Hritonenko V, Mun JJ, Tam C, Simon NC, Barbieri JT, Evans DJ, Fleiszig SM: Adenylate cyclase activity of Pseudomonas aeruginosa ExoY can mediate bleb-niche formation in epithelial cells and contributes to virulence. Microb Pathog 2011, 51:305–12.

[17] Jolly AL, Takawira D, Oke OO, Whiteside SA, Chang SW, Wen ER, Quach K, Evans DJ, Fleiszig SM: Pseudomonas aeruginosa-induced bleb-niche formation in epithelial cells is independent of actinomyosin contraction and enhanced by loss of cystic fibrosis transmembrane-conductance regulator osmoregulatory function. mBio 2015, 6:e02533.

[18] Couch Y, Buzàs EI, Di Vizio D, Gho YS, Harrison P, Hill AF, Lötvall J, Raposo G, Stahl PD, Théry C, Witwer KW, Carter DRF: A brief history of nearly EV-erything - The rise and rise of extracellular vesicles. J Extracell Vesicles 2021, 10:e12144.

[19] Tkach M, Théry C: Communication by Extracellular Vesicles: Where We Are and Where We Need to Go. Cell 2016, 164:1226–32.

[20] Ayilam Ramachandran R, Lemoff A, Robertson DM: Pseudomonas aeruginosa-Derived Extracellular Vesicles Modulate Corneal Inflammation: Role in Microbial Keratitis? Infect Immun 2023, 91:e0003623.

[21] Ayilam Ramachandran R, Lemoff A, Robertson DM: Extracellular vesicles released by host epithelial cells during Pseudomonas aeruginosa infection function as homing beacons for neutrophils. Cell Commun Signal 2024, 22:341.

[22] Ayilam Ramachandran R, Abdallah JT, Rehman M, Baniasadi H, Blanton AM, Vizcaino S, Robertson DM: Pseudmonas aeruginosa impairs mitochondrial function and metabolism during infection of corneal epithelial cells. bioRxiv 2024.

[23] Robertson DM, Li L, Fisher S, Pearce VP, Shay JW, Wright WE, Cavanagh HD, Jester JV: Characterization of Growth and Differentiation in a Telomerase-Immortalized Human Corneal Epithelial Cell Line. Invest Ophthalmol Vis Sci 2005, 46:470–8.

[24] Yuan M, Breitkopf SB, Yang X, Asara JM: A positive/negative ion–switching, targeted mass spectrometry–based metabolomics platform for bodily fluids, cells, and fresh and fixed tissue. Nature Protocols 2012, 7:872–81.

[25] Al-Bakheit A, Traka M, Saha S, Mithen R, Melchini A: Accumulation of Palmitoylcarnitine and Its Effect on Pro-Inflammatory Pathways and Calcium Influx in Prostate Cancer. Prostate 2016, 76:1326–37.

[26] Rada B: Interactions between Neutrophils and Pseudomonas aeruginosa in Cystic Fibrosis. Pathogens 2017, 6.

[27] Mun Y, Hwang JS, Shin YJ: Role of Neutrophils on the Ocular Surface. Int J Mol Sci 2021, 22.

[28] Finkel T, Menazza S, Holmström KM, Parks RJ, Liu J, Sun J, Liu J, Pan X, Murphy E: The ins and outs of mitochondrial calcium. Circ Res 2015, 116:1810–9.

[29] Meadows JA, Willsey GG, Wargo MJ: Differential requirements for processing and transport of short-chain versus long-chain O-acylcarnitines in Pseudomonas aeruginosa. Microbiology (Reading) 2018, 164:635–45.

[30] Meadows JA, Wargo MJ: Carnitine in bacterial physiology and metabolism. Microbiology (Reading) 2015, 161:1161–74.

[31] Meadows JA, Wargo MJ: Characterization of Pseudomonas aeruginosa growth on O-acylcarnitines and identification of a short-chain acylcarnitine hydrolase. Appl Environ Microbiol 2013, 79:3355–63.

[32] Barnabei L, Laplantine E, Mbongo W, Rieux-Laucat F, Weil R: NF-κB: At the Borders of Autoimmunity and Inflammation. Front Immunol 2021, 12:716469.

[33] Guichard C, Pedruzzi E, Dewas C, Fay M, Pouzet C, Bens M, Vandewalle A, Ogier-Denis E, Gougerot-Pocidalo MA, Elbim C: Interleukin-8-induced priming of neutrophil oxidative burst requires sequential recruitment of NADPH oxidase components into lipid rafts. J Biol Chem 2005, 280:37021–32.

[34] Bogdan ED, Stuard WL, Titone R, Robertson DM: IGFBP-3 Mediates Metabolic Homeostasis During Hyperosmolar Stress in the Corneal Epithelium. Invest Ophthalmol Vis Sci 2021, 62:11.

[35] Stuard WL, Guner MK, Robertson DM: IGFBP-3 Regulates Mitochondrial Hyperfusion and Metabolic Activity in Ocular Surface Epithelia during Hyperosmolar Stress. Int J Mol Sci 2022, 23.

[36] Mussi N, Stuard WL, Sanches JM, Robertson DM: Chronic Hyperglycemia Compromises Mitochondrial Function in Corneal Epithelial Cells: Implications for the Diabetic Cornea. Cells 2022, 11.

