## Supplementary figures for "*Pseudomonas aeruginosa* infection increases palmitoyl carnitine release by host-derived extracellular vesicles"

**Supplementary figure 1:** Mass spectrometry proteomic analysis confirmed a reduction in expression of carnitine transferase enzymes in PA-C EVs. (A) Carnitine O-palmitoyl transferase 2, and (B) carnitine O-acetyl transferase. The peak areas detected by the mass spectrometer were normalized using total protein for comparison between the two groups. Data represented as mean ± standard deviation, N=3, t-test, *p<0.05.

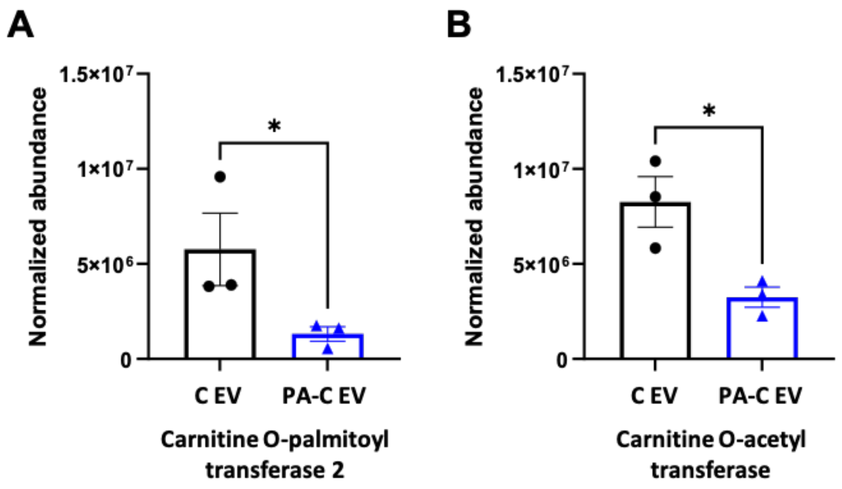

**Supplementary figure 2: 100 µM PAMC induced toxicity in hTCEpi cells.** (A-C) A lactate dehydrogenase assay was used to assess PAMC toxicity. Cells were assessed after 24 hours (A), 48 hours (B), and 72 hours. Data is represented as mean ± standard deviation, N≥3, one-way ANOVA with Tukey's post hoc multiple comparison test, **p<0.01, ****p<0.0001. VC: vehicle control (DMSO), NTC: no template control.

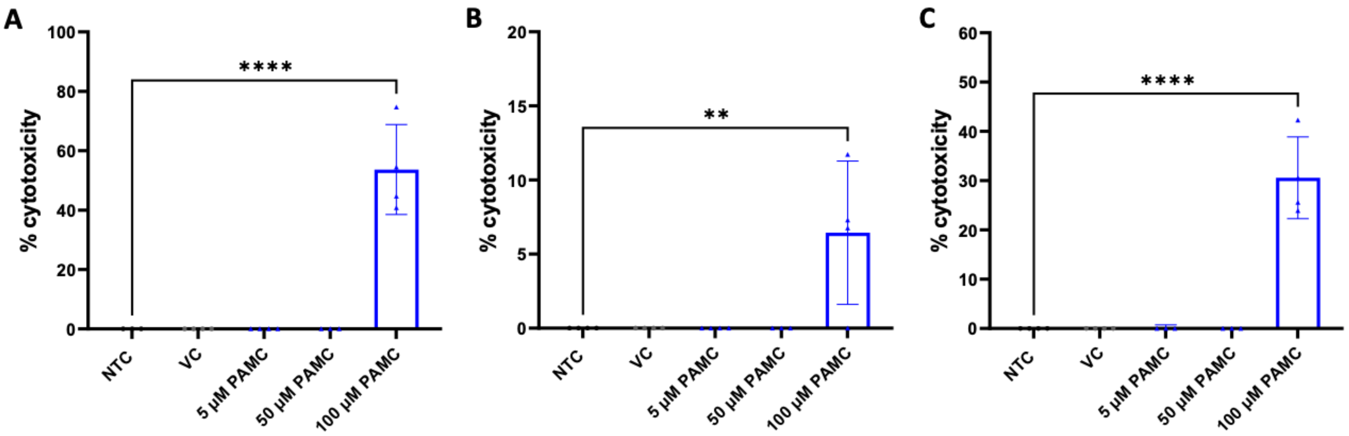

**Supplementary figure 3: Two hours of pre-treatment was sufficient to reduce intracellular levels of PA in hTCEpi cells.** The cells were treated with palmitoyl carnitine at a concentration 50 µM for 2 hours prior to inoculated. A 1 hour gentamicin protection assay was used to detect intracellular levels of PA. Mean ± standard deviation, N=3, one-way ANOVA with Tukey's post hoc multiple comparison test, *<p0.05, **p<0.01.

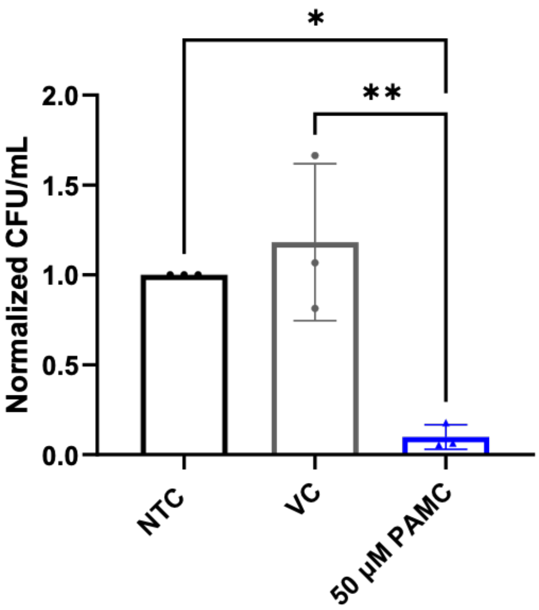
